# A genetic and physiological model of renal dysfunction in Lowe syndrome

**DOI:** 10.1101/2024.01.15.575703

**Authors:** Navyashree A Ramesh, Vaishali Kataria, Indra Sara Lama, Rajan Thakur, Avishek Ghosh, Sanjeev Sharma, Aishwarya Venugopal, Anil Vasudevan, Raghu Padinjat

**Affiliations:** National Centre for Biological Sciences, TIFR-GKVK Campus, Bellary Road, Bengaluru-560065, India

**Author notes:** Department of Pediatric Nephrology, St. John’s Medical College Hospital, Bengaluru-560034, India. Department of Neuroscience, Brown University, RI 02906, USA. Molecular, Cellular, and Developmental Biology, University of Michigan, 2220 Biological Science Building, 1105 North University Ave., Ann Arbor, MI 48109-1085, USA. Vascular Biology Program, Boston Children’s Hospital,1 Blackfan Street, Karp Research Building, 12th Floor, Boston, MA 02115, USA. Depts. of Biochemistry, Weill-Cornell Medicine,1300 York Avenue, A403-412, New York, NY 10065, USA.

## Abstract

Lowe syndrome (LS) is an X-linked recessive genetic disorder characterized by renal dysfunction, neurodevelopmental defects, and cataract. The affected gene, *OCRL* encodes for a polyphosphoinositide 5-phosphatase. OCRL is localized to multiple sub-cellular locations in the endolysosomal system and defects in these organelles have been described in human cells depleted of OCRL. However, the relationship of the endolysosomal defects in OCRL depleted cells to the altered physiology of kidney cells of LS patients has not been completely determined. Here we model the kidney phenotypes of LS using a *Drosophila* nephrocyte model. Using this model system, we demonstrate that OCRL plays a cell-autonomous role in nephrocyte function. Deletion of the only OCRL ortholog in *Drosophila* (*dOCRL*) leads to cell-autonomous defects in larval nephrocyte structure and function. Null mutants of *dOCRL* (*dOCRL^KO^*) show defects in the endolysosomal system of larval nephrocytes that are associated with physiological defects in nephrocyte function. These defects could be rescued by reconstitution with a human *OCRL* transgene but not with a phosphatase dead version or a human LS patient derived mutation. Overall, this work provides a model system to understand the mechanisms by which the sub-cellular changes from loss of OCRL leads to defects in kidney function in human patients.

## Introduction

Lowe syndrome (LS) which is also known as oculocerebrorenal syndrome of Lowe is a rare X-linked recessive genetic disorder characterized by clinical features of renal tubular dysfunction, mental retardation, and cataract (De Matteis et al., 2017; Preston et al., 2020). The disease results from the mutations in the *OCRL* gene that encodes a type II family inositol polyphosphate 5-phosphatase enzyme involved in the hydrolysis of phosphate at 5^th^ position of phosphatidylinositol-4,5-bisphosphate [PI(4,5)P_2_] to generate PI4P(Attree et al., 1992; Zhang et al., 1995). The *OCRL* gene is widely expressed across human cells and tissues and throughout development. OCRL is a multi-domain enzyme with a catalytic 5-phosphatase domain, N-terminal pleckstrin homology (PH) domain, an ASH domain and a RhoGAP domain (De Matteis et al., 2017). The OCRL protein is localized to multiple sub-cellular membranes in human cells including the plasma membrane, endocytic compartments, trans-Golgi, cilium and lysosomes [reviewed in (Mehta et al., 2014)] and the additional non-phosphatase domains of OCRL are thought to help localize the protein to distinct sub-cellular locations.

Proximal tubular dysfunction is a key clinical manifestation of LS which is characterized by renal tubular acidosis, low molecular weight proteinuria, hypercalciuria, albuminuria, aminoaciduria, carnitine wasting, phosphaturia with perturbed glomerular filtrate rate that deteriorates with age (Bockenhauer et al., 2008; Recker et al., 2015). Several studies have attempted to decipher the sub-cellular basis of altered proximal tubular function in LS. Studies in COS-7 (kidney derived) cells found OCRL localized to early endosomes and the Golgi apparatus (Ungewickell et al., 2004) and early endosomal function was impaired by OCRL depletion(Vicinanza et al., 2011). Other studies with human kidney biopsy samples showed OCRL localized on lysosomes (Zhang et al., 1998) and a role for OCRL in regulating autophagosome-lysosomal function has been proposed (De Leo et al., 2016). Studies in a zebrafish model of LS have reported alterations in the endosomal system (Oltrabella et al., 2015) and this has also been reported in a humanized mouse model of LS (Festa et al., 2019). Despite these sub-cellular defects that have been described on OCRL depletion, the relevance of any of these to altered renal physiology has not been established.

The filtration of blood in the glomerulus of nephrons followed by selective reabsorption of essential molecules is a key aspect of human kidney function. In *Drosophila* too, nephrocytes perform an analogous function to purify circulating hemolymph. *Drosophila* larvae have two types of nephrocytes garland and pericardial, tethered to the esophagus and heart respectively. Nephrocytes uptake hemolymph by endocytosis, sort the endocytosed cargo for degradation in lysosomes or recycle it back to the cell surface. This endocytic function is like that seen in proximal tubular cells and hence offers avenues to model various aspects of proximal tubular functions (Helmstädter et al., 2017) (Koehler and Huber, 2023). Owing to their large size, nephrocytes allow both live cell imaging of organelle structure, function, and analysis of physiological readouts from intact animals. In contrast to human genome, the *Drosophila* genome contains a single homolog of *OCRL* (Balakrishnan et al., 2015; Ben El Kadhi et al., 2011) and therefore offers an opportunity to analyze function without the complexity of contribution from multiple genes that encode the same activity.

In this study, we have developed an *in vivo Drosophila* model of LS with a goal of understanding the mechanisms that may underlie the renal dysfunction in these patients. We generated a whole-body null allele of *dOCRL* (*dOCRL^KO^)* that was late larval to pupal lethal with reduced body mass, delayed growth and development of larvae. Using larval pericardial nephrocytes as a model, we find multiple defects in the endolysosomal system in *dOCRL^KO^* along with a physiological defect in clearing silver nitrate, a heavy metal toxin through the nephrocytes. Using nephrocyte specific deletion of dOCRL, we find that these defects are cell autonomous to this cell type. Lastly, we were able to rescue these phenotypes with a human OCRL transgenic but not on carrying a patient-specific mutation in OCRL. Thus, our findings provide an animal model which can be used to investigate the functional patient mutations in OCRL and understand the mechanistic basis of proximal tubular dysfunction in these patients.

## Results

### *dOCRL* is required for growth and development of the larvae

The *Drosophila* genome encodes a single ortholog of human *OCRL* named as *dOCRL* (CG3573) that is located on the X chromosome. We have previously generated a null mutant of *dOCRL* (*dOCRL^KO^)* using CRISPR/Cas9 genome engineering (Trivedi et al., 2020). Quantitative RT-PCR analysis showed virtually no *dOCRL* transcript in *dOCRL^KO^* animals (Fig 1A) and a western blot with antibodies specific to the protein showed no detectable product (Fig 1G). We observed that homozygous *dOCRL^KO^* larvae were lethal at the second instar stage when grown on standard fly food, but when grown on yeast medium, they reach the 3^rd^ larval instar or pupal stage, but no adult flies eclosed. *dOCRL^KO^* larvae showed lethality at every stage of development when compared to wild type (Fig 1B). When *dOCRL^KO^* was grown on yeast medium, the animals showed a developmental delay of ca. 43 hours (h) for 50% of animals to pupariation (Fig 1C) and only about 58% of larvae pupariated. In addition, wandering larvae were smaller and weighed only about 50% of the body weight of controls (Fig 1D,1F). Both phenotypes were rescued by the transgenic expression of a *dOCRL* using *hs-GAL4* which is expressed ubiquitously (Fig 1D-F). However, the adult lethality was not rescued, presumably due to insufficient expression of dOCRL with *hs-Gal4*. Lethality was however completely rescued (data not shown) by reconstitution using an X-duplication (BL-31454), which includes *dOCRL* genomic region. Collectively, these results suggest that *dOCRL* is required for normal growth and larval development in *Drosophila*.

**Figure 1:**
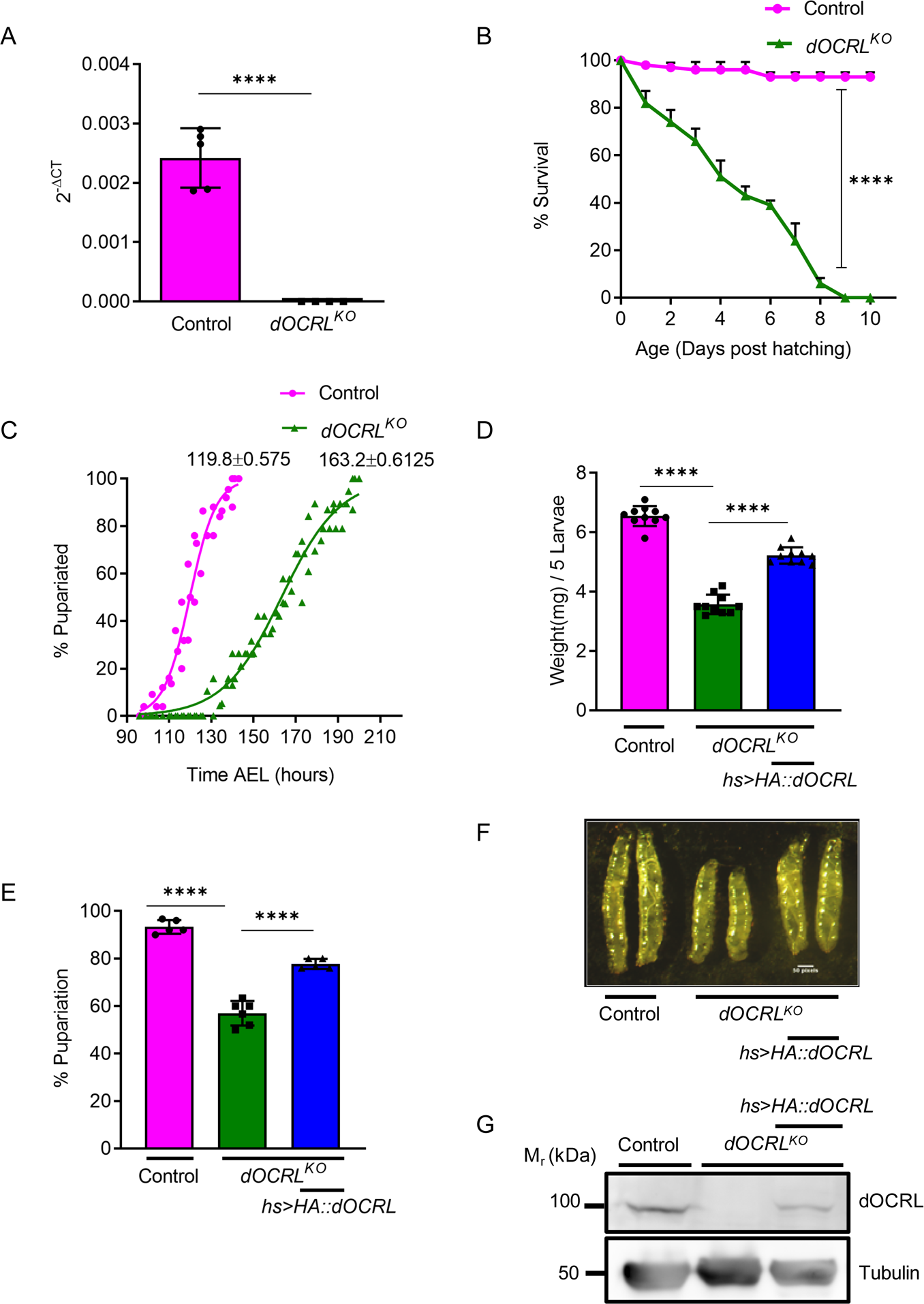
dOCRL regulates growth and development of larvae. (A) Quantitative RT-PCR to check transcript levels of *dOCRL* in Control and *dOCRL^KO^*. (B) Survival plot of *dOCRL^KO^* and control larvae, Y axis represents the percentage of survival of larvae and X axis represents the days post larval hatching. (C) represents the growth curve to analyze the percentage of pupariation over time after egg laying (AEL) for both control and *dOCRL^KO^*. (D) Average weight of third instar wandering larvae of control, *dOCRL^KO^* and *dOCRL^KO^* reconstituted with the wild type *dOCRL* gene (*dOCRL^KO^;hs>HA::dOCRL*) was calculated. (E) Percentage of pupariation was calculated for the wild type, *dOCRL^KO^* and *dOCRL^KO^;hs>HA::dOCRL*. (F) Micrographs of wild type, *dOCRL^KO^* and *dOCRL^KO^;hs>HA::dOCRL* at third instar wandering stage. We observed that *dOCRL^KO^* is lethal at pupal stage/late 3^rd^ instar. (G) Immunoblots for verification of dOCRL protein expression in control, *dOCRL^KO^* and *dOCRL^KO^;hs>HA::dOCRL*. Statistical tests: (A, D, E) Column plots with mean ± S.E.M are shown. Two tailed unpaired t-test with welch correction is used. ****p<0.0001, ***p<0.001, **p<0.01. (B) Survival curve used Long-rank (Mantel-cox) test and Gehan-Breslow-Wilcoxon test to analyze the statistical significance between two genotypes, ****p<0.0001.

### *dOCRL* regulates PI(4,5)P_2_ and PI4P in nephrocytes

OCRL is proposed to dephosphorylate PI(4,5)P_2_ to generate PI4P and thus maintain the balance of these two lipids. We quantified the levels of PI(4,5)P_2_ at the plasma membrane of pericardial nephrocytes. For this, we used the PH domain of PLCδ tagged to mCherry (PH-PLCδ::mCherry), a probe that specifically binds to PI(4,5)P_2_ (Hammond and Balla, 2015). We observed that in wild type nephrocytes, PH-PLCδ::mCherry uniformly decorated the plasma membrane with some punctate structures just below the plasma membrane (Fig 2A). In *dOCRL^KO^* nephrocytes PH-PLCδ::mCherry accumulated at higher levels on the plasma membrane with many punctate structures in the cytoplasm just below the plasma membrane (Fig 2A). The levels of PI(4,5)P_2_ at the plasma membrane were quantified by estimating the ratio of plasma membrane/cytoplasmic fluorescence and was higher in *dOCRL^KO^* compared to controls (Fig 2 B). Likewise, using the P4M::GFP probe that binds to PI4P (Balakrishnan et al., 2018) and mainly decorates the plasma membrane of nephrocytes (Fig 2 C), we estimated changes in PI4P levels. Using this approach, we found that the PI4P level at the plasma membrane was lower in *dOCRL^KO^* compared to controls (Fig 2D). Importantly, both the PH-PLCδ::mCherry and P4M::GFP proteins were expressed equally in wild type and *dOCRL^KO^* (Sup Fig1 A-D). We also measured total PIP and PIP_2_ levels from whole larval lipid extracts using liquid chromatography coupled to tandem mass spectrometry (LC-MS/MS). We observed a trend of increased total PIP_2_ between *dOCRL^KO^* and controls, however total PIP levels were unchanged (Fig 2E, F). Expressing the *dOCRL* transgene in *dOCRL^KO^* could reduce PIP_2_ levels but did not alter total PIPs (Fig 2E, F).

**Figure 2:**
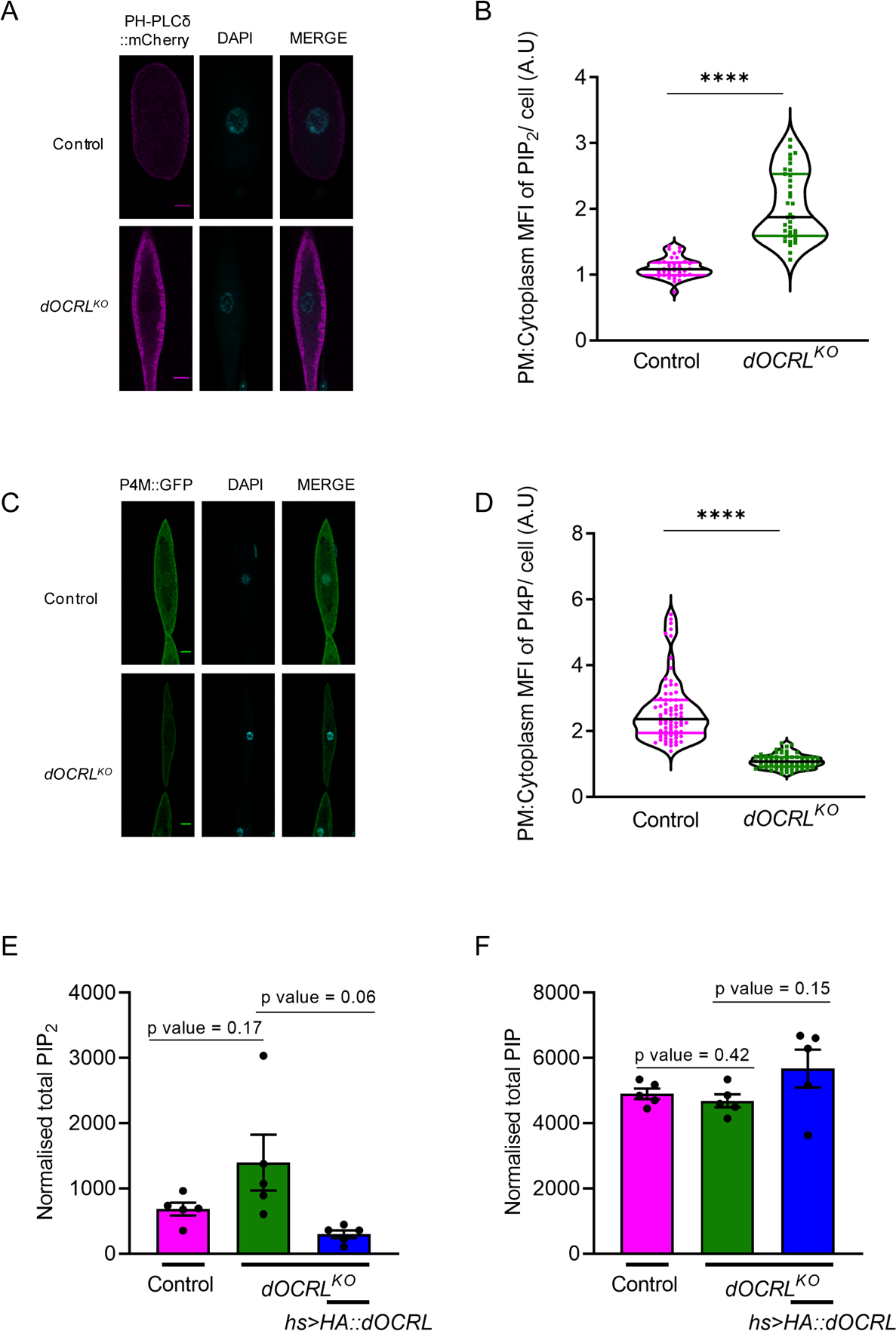
dOCRL is required to maintain the levels of PIP_2_ and PI4P in nephrocytes. (A-B) Confocal micrographs and quantification of PI(4,5)P_2_ in control and *dOCRL^KO^* was done using PH-PLCδ::mCherry probe. (C-D) PI4P was measured similarly using P4M::GFP as a probe. (E-F) Whole larval PIP and PIP_2_ levels were quantified by LC-MS/MS. Statistical tests: (B, D) Column graph is plotted with individual dot representing the value of each nephrocytes with mean ± S.E.M are shown. Two tailed unpaired t-test is used. ****p<0.0001, ***p<0.001, **p<0.01. (E,F) Unpaired t-test with Welch’s correction is used ****p<0.05.

### *dOCRL* is required for nephrocyte function in *Drosophila*

To investigate the filtration function of nephrocytes we measured the clearance of the heavy metal silver, using silver nitrate (AgNO_3_) from these cells. For this, we transferred first instar larvae to feed on 0.003% AgNO_3_ in yeast paste and after 32 h, these larvae were transferred to yeast paste without AgNO_3_. Subsequently, larvae were removed from the AgNO_3_ free medium at defined time points, dissected and the amount of AgNO_3_ in each nephrocyte visualized and quantified (Fig 3A). This analysis was done on larvae at 36, 42, and 52 h post transfer from medium containing AgNO_3_ onto non-AgNO_3_ medium. Nephrocytes were imaged under bright field microscope and these images were converted to 8-bit images (16 color pixels) corresponding to the intensity of AgNO_3_, from white being highest concentration to black being least concentration. It was observed that in wild type animals, there were only traces of AgNO_3_ remaining by 36 h post-transfer but at 52 h AgNO_3_ was completely cleared out (Fig 3B). However, in *dOCRL^KO^* nephrocytes, AgNO_3_ was not cleared at 36, 42, and 52 h post-transfer and a large proportion of nephrocytes containing AgNO_3_ could be observed (Fig 3B) and this clearance defect could be rescued by reconstituting with the wild type *dOCRL* (Fig 3B). We quantified the percentage of cells containing varying amounts of AgNO_3._ The micrographs were converted into 16 color pixels and the cells with high (red) and low (yellow) levels of AgNO_3_ were counted. For each genotype we then counted the proportion of cells with red and yellow pixels. The results, represented in Fig 3C, show that at all time points studied, *dOCRL^KO^* had a significantly higher percentage of cells with red pixels compared to wild type; this phenotype was rescued by reconstitution of *dOCRL^KO^* with wild type *dOCRL* (Fig 3C). These findings demonstrate that *dOCRL* function is required for AgNO_3_ clearance in *Drosophila* nephrocytes.

**Figure 3:**
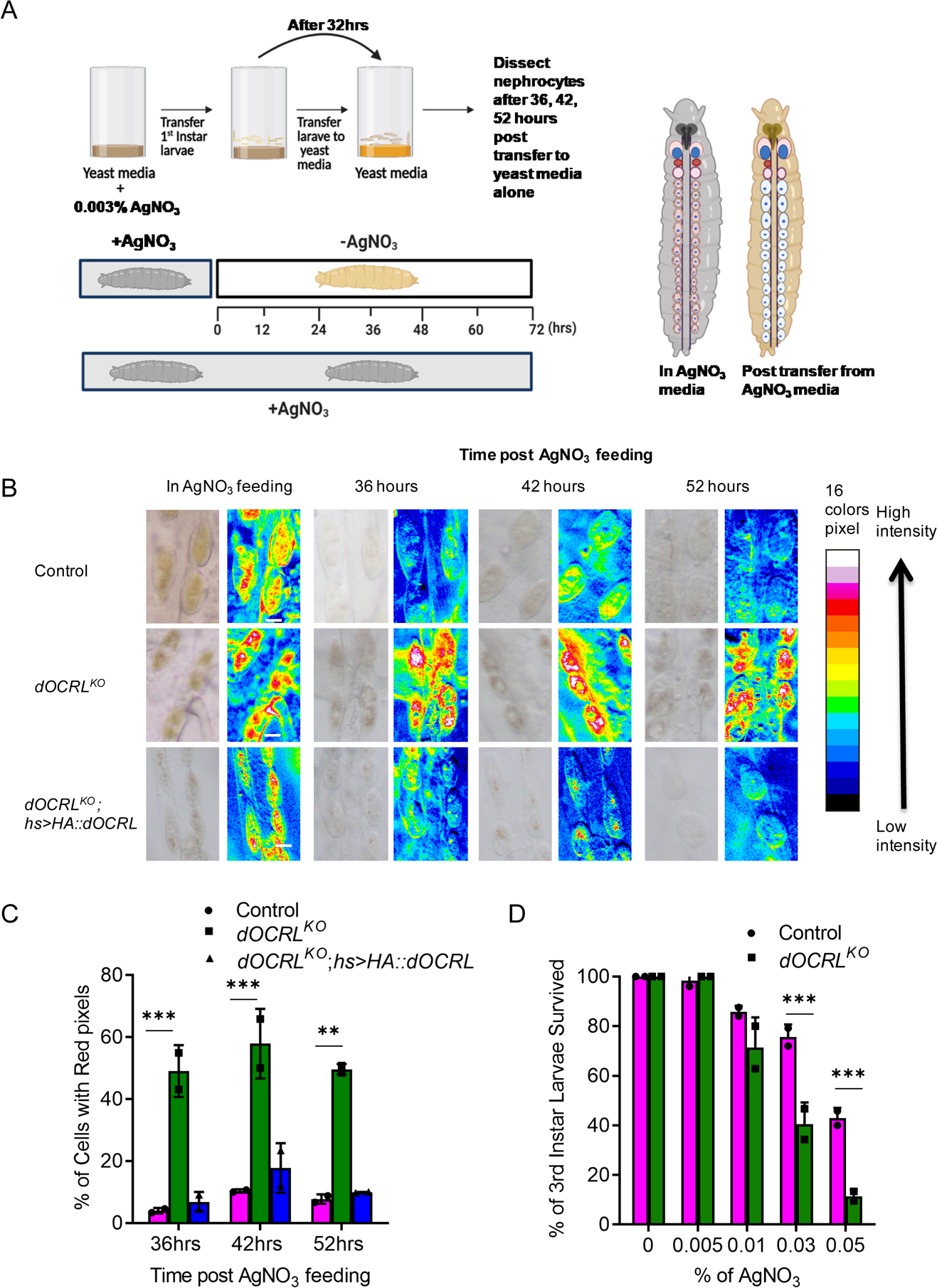
Loss of *dOCRL* perturbs filtration function in nephrocytes. (A) A cartoon illustrating the method of silver nitrate AgNO_3_ clearance assay. (B) Micrographs of dissected nephrocytes from control, *dOCRL^KO^*, and *dOCRL^KO^;hs>HA::dOCRL* larvae showing AgNO_3_ clearance at 36,42 and 56 hours after transfer from AgNO_3_ yeast media to just yeast media and their respective 8-bit 16 color pixel converted micrographs. 16 color pixels are arranged in the order of intensity levels from minimum with black to maximum level with white. (C) Percentage of cells with red pixel intensity corresponding to highest levels of AgNO_3_ in nephrocytes of control, *dOCRL^KO^*, *dOCRL^KO^;hs>HA::dOCRL* at 36,42 and 56 hours after transfer. (D) Survival of larvae til third instar grown on increasing concentrations of AgNO_3._ Statistical tests: (C, D) XY plots with mean ± S.E.M are represented. Two-way ANOVA grouped analysis with Bonferroni’s post multiple comparison tests was performed using graph pad prism to compare between each group. Ns-Non significance, ****p<0.0001, ***p<0.001, *p<0.05. This statistical significance is represented on graph.

It has been reported that clearance of AgNO_3_ by nephrocytes is essential for larval survival when grown on food containing this heavy metal, and if nephrocyte function is perturbed, feeding with AgNO_3_ leads to larval lethality (Ivy et al., 2015). Since we noted slower clearance of AgNO_3_ in *dOCRL^KO^*, we tested the sensitivity of wild type and *dOCRL^KO^*to growth on AgNO_3_. Newly hatched first instar larvae were transferred onto fresh medium containing defined concentrations of AgNO_3_. The percentage of larvae that survive to the third larval instar were plotted at each AgNO_3_ concentration to that in control media. In wild type larvae, lethality was observed when grown at 0.01% AgNO_3_ or higher with lethality increasing with increasing AgNO_3_ concentration (Fig 3D). In *dOCRL^KO^* we observed higher percentage of larval lethality at 0.03% and 0.05% AgNO_3_ as compared to controls (Fig 3D). These results suggest that *dOCRL* is required to protect *Drosophila* larvae against heavy metal induced larval lethality.

### *dOCRL* is required for endocytosis in nephrocytes

The endocytosis of fluid filtered through the diaphragm is a key part of nephrocyte function and it has been reported that nephrocytes internalize dextran by fluid phase endocytosis (Grawe et al., 2009). Since OCRL has been implicated in regulating endocytosis in cultured mammalian cells,we studied this process using 10 kDa TMR-Dextran as a cargo in an *ex vivo* endocytosis assay. As a negative control, uptake assays were performed at 4^0^C since endocytosis is blocked at this temperature. At 25 ^0^C robust endocytosis of TMR-Dextran was seen which was blocked at 4^0^C (Fig 4A, B). By contrast in *dOCRL^KO^* there was complete absence of TMR-Dextran uptake at 25 ^0^C (Fig 4A, B). This defect in uptake could be rescued by reconstitution of *dOCRL^KO^* with *hs*>*dOCRL* (Fig 4 A, B). We also assessed clathrin-mediated endocytosis in nephrocytes by using maleic anhydride-BSA (mBSA) conjugated to Cy5 bis functional dye (Abrams et al., 1992). In *dOCRL^KO^*nephrocytes, mBSA uptake was significantly reduced at 25^0^C compared to controls and this was rescued when we reconstituted *dOCRL^KO^*with hs>*dOCRL* (Fig 4 C, D). These results demonstrate that *dOCRL* is required for both fluid phase and clathrin dependent endocytosis.

**Figure 4:**
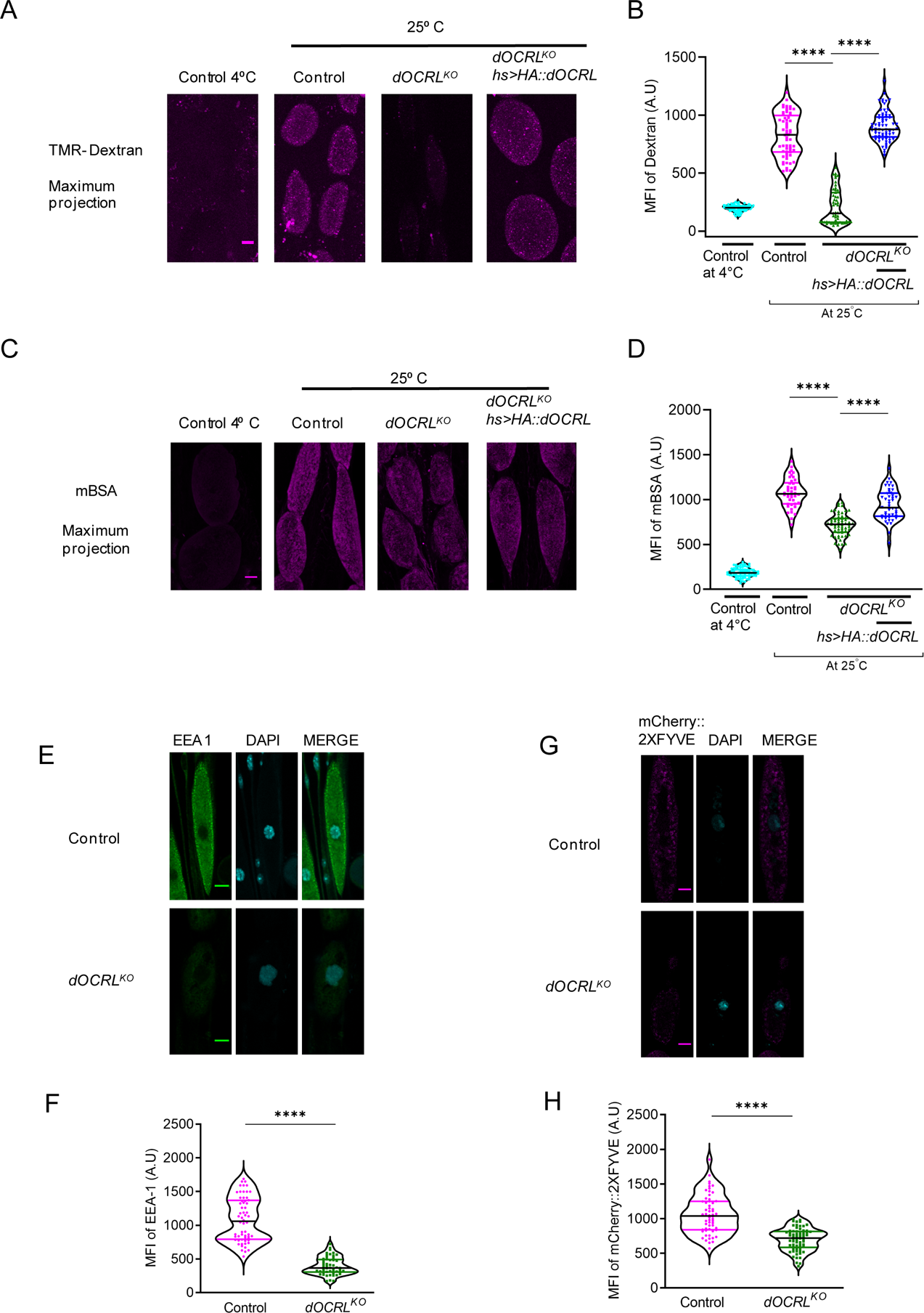
Loss of dOCRL perturbs the endocytosis in nephrocytes. (A) Confocal micrographs of 10KDa TMR-dextran uptake assay in control, *dOCRL^KO^* and *dOCRL^KO^;hs>HA::dOCRL.* (B) Mean fluorescence intensity (MFI) of 10KDa TMR-Dextran in nephrocytes per unit area calculated using imageJ. (C) Confocal micrographs of mBSA-Cy5 uptake assay in control, *dOCRL^KO^*and *dOCRL^KO^;hs>HA::dOCRL.* (D) Quantification of uptake of mBSA by calculating the MFI. (E-F) Confocal micrographs and quantification of Immunostaining of control, *dOCRL^KO^* nephrocytes with early endosome marker EEA-1, nucleus of these cells was stained with DAPI. (G) Confocal micrographs of nephrocytes labelled with 2XFYVE domain. UAS-mCherry::2XFYVE is driven in nephrocytes using Dot Gal4. (H) Quantification of MFI of mCherry::2XFYVE by Imagej. Statistical tests: XY plots with mean ± S.E.M are represented. Two tailed unpaired t-test is used. ****p<0.0001, ***p<0.001, **p<0.01.

### Loss of *dOCRL* perturbs endosomal compartments in nephrocytes

Given the endocytic defects that we observed, we visualized endosomal compartments in *dOCRL^KO^* nephrocytes. We performed immunostaining with EEA-1, a marker of early endosomes and found that EEA-1 staining was substantially reduced in *dOCRL^KO^* compared to controls (Fig 4E, F). The lipid phosphatidylinositol 3-phosphate (PI3P) is enriched on early endosomes and can be visualized using the 2XFYVE domain fused to a fluorescent protein (mCherry::2XFYVE); we found that the intensity of mCherry::2XFYVE punctae was lower in *dOCRL^KO^* compared to controls (Fig 4G, H). Lastly, we also observed that the intensity of Rab7 punctae, marking late endosomes, was lower in *dOCRL^KO^* (Fig 5A, B). This suggests that in nephrocytes, depletion of dOCRL impacts the early and late endocytic system.

**Figure 5:**
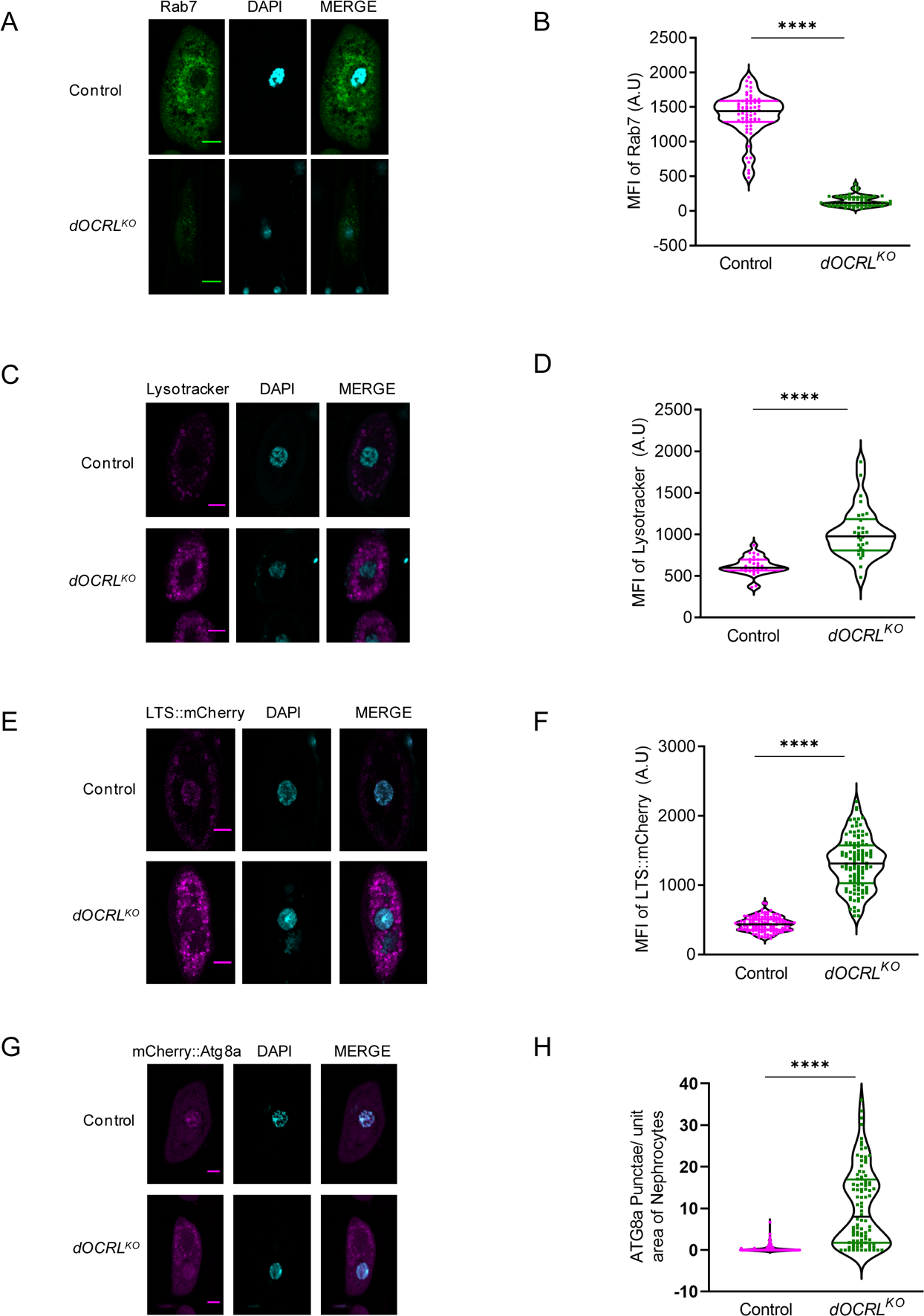
*dOCRL^KO^* is required to maintain organelle homeostasis. (A) Confocal micrographs of Immunostaining of control and *dOCRL^KO^*nephrocytes with late endosome marker Rab7,(B) Mean fluorescence intensity (MFI) of Rab7 antibody staining calculated using imagej. (C) Confocal micrographs of nephrocytes stained with lysotracker deep red to label the acidified compartments of control and *dOCRL^KO^*. (D) MFI of Lysotracker staining measured using ImageJ. (E) Confocal micrograph of nephrocytes labelled with lysosome targeting probe LTS::mcherry in control and *dOCRL^KO^* (F) MFI of Lysosome levels in nephrocytes measured using imagej. (G) Confocal micrograph of nephrocytes labelled with autophagosomes targeting probe mcherry::Atg8a in control and *dOCRL^KO^*.(H) Number of Atg8a puncta in nephrocytes in control and *dOCRL^KO^* were counted using ImageJ and normalized to unit area of nephrocytes. Statistical tests: XY plots with mean ± S.E.M are represented. Two tailed unpaired t-test is used. ****p<0.0001, ***p<0.001, **p<0.01.

### dOCRL depletion alters cellular degradative organelles

We investigated the levels of lysosomes, acidified vesicles in nephrocytes by staining with lysotracker that reports acidified subcellular compartments. We observed elevated lysotracker staining in *dOCRL^KO^* nephrocytes compared to wild type. We quantified the fluorescence intensity of lysotracker staining and found that *dOCRL^KO^* nephrocytes showed significantly increased levels of lysotracker (Fig 5 C, D).

We analyzed if the enhanced lysotracker staining was due to increased lysosomes. To visualize lysosomes we developed a construct in which the lysosome targeting sequence (LTS) is fused to mCherry (LTS::mCherry) (Ghosh et al., 2023). When expressed in control nephrocytes, LTS::mCherry marks punctate vesicular structures that can be quantified using confocal microscopy. We noted that the mean fluorescence intensity of LTS::mCherry was substantially elevated in *dOCRL^KO^* nephrocytes compared to control (Fig 5 E, F). A sub-population of lysosomes come from autophagosome-lysosome fusion; therefore, we looked at the levels of autophagosomes by expressing mCherry::Atg8a in nephrocytes. We observed significant increase in the number of mCherry::Atg8a punctae in *dOCRL^KO^* nephrocytes compared to controls (Fig 5 G, H).

### Cell autonomous regulation of nephrocyte function by dOCRL

*dOCRL^KO^* larvae show a whole-body growth phenotype along with defects in nephrocyte structure and function. To test if the requirement for *dOCRL* to support nephrocyte function is cell-autonomous, we generated a nephrocyte specific knockout of *dOCRL* using the CRISPR/Cas9 gene editing by expressing Cas9 only in nephrocytes using *Dot GAL4* (Trivedi et al., 2020). We obtained an amplicon of 555 bp confirming the deletion of *dOCRL* in nephrocytes (Sup Fig 2A); we refer to this allele as *dOCRL^N-KO^*. Using the PH-PLCδ::mCherry probe, we found that PIP_2_ levels in *dOCRL^N-KO^* were significantly increased as compared to the control (Fig 6A, Sup Fig 2E, F).

**Figure 6:**
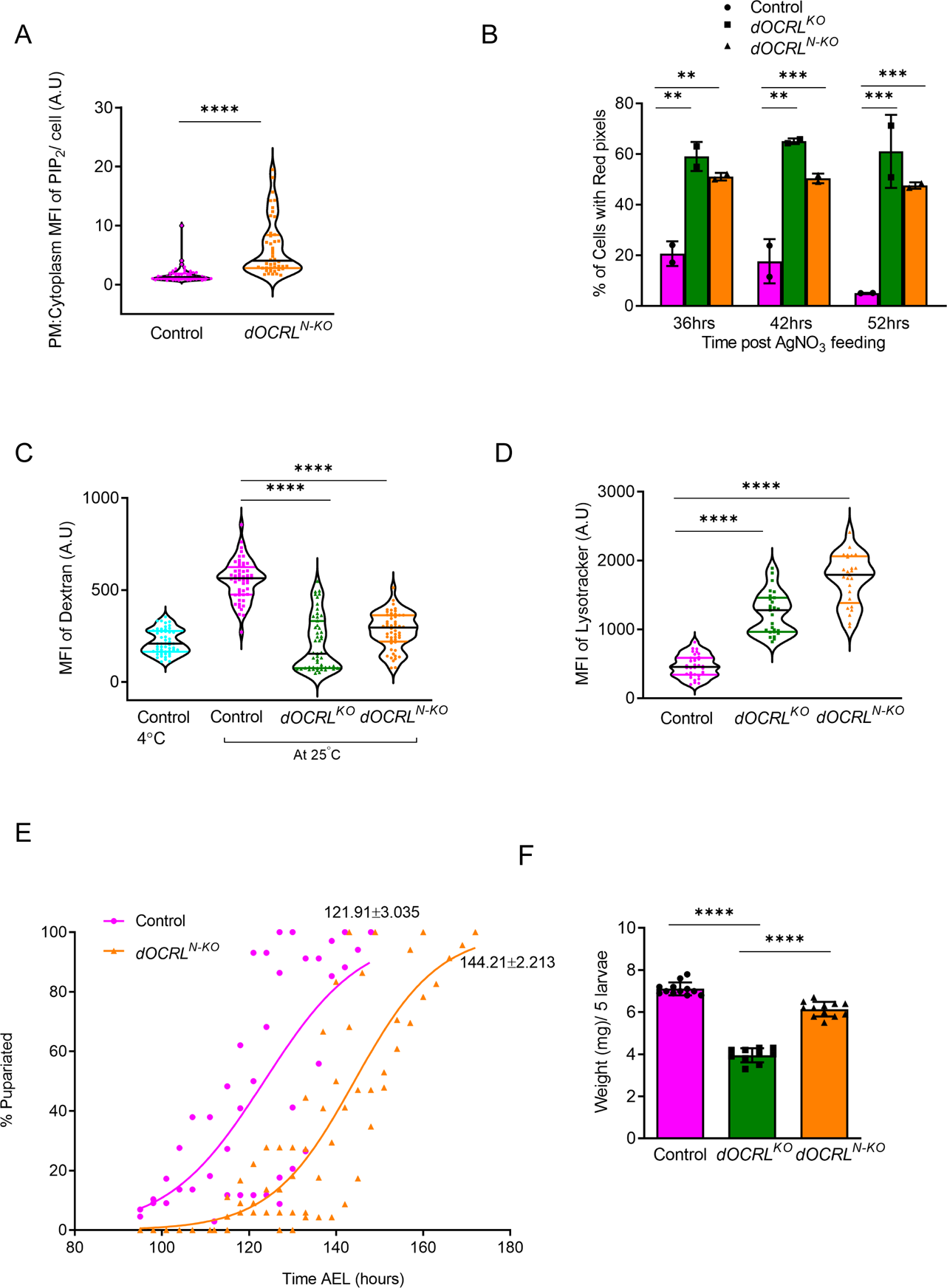
Generation of Nephrocyte specific *dOCRL^N-KO^*. (A) Quantification of plasma membrane to cytoplasmic MFI levels of PIP_2_ in control and *dOCRL^N-KO^.* (B) Percentage of cells with red pixel intensity corresponding to highest levels of AgNO_3_ in nephrocytes at 36h, 42h, and 56h after transfer. (C) Quantification of mean fluorescence intensity (MFI) of 10KDa TMR-Dextran in nephrocytes calculated using imageJ. (D) Quantification of MFI of Lysotracker staining in nephrocytes of control and *dOCRL^N-KO^* measured using imageJ (E) Growth curve to analyze the percentage pupariation over time after egg laying (AEL) for both control and *dOCRL^N-KO^*. (F) Average weight of third instar wandering larvae of control, *dOCRL^KO^,dOCRL^N-KO^* was measured and plotted. Each data set corresponds to weight in milligram per 5 larvae. Statistical tests: XY plots with mean ± S.E.M are represented. (A, B, E, F) Two tailed unpaired t-test is used. ns-Non significance, ****p<0.0001, ***p<0.001, **p<0.01. (D) Two-way ANOVA grouped analysis with Bonferroni’s post multiple comparison tests was performed using graph pad prism to compare between each group. ns-Non significance, ****p<0.0001, ***p<0.001, *p<0.05. This statistical significance is represented on graph.

We estimated nephrocyte function in *dOCRL^N-KO^* using the AgNO_3_ clearance assay. In control nephrocytes, by 36 h post transfer onto normal medium, AgNO_3_ was seen in only 20% of nephrocytes. By contrast, in *dOCRL^N-KO^*, ca. 60% of nephrocytes showed the presence of AgNO_3_, recapitulating that seen in the whole animal *dOCRL^KO^* (Fig 6B). Likewise, we also measured fluid phase endocytosis in *dOCRL^N-KO^* and found reduced TMR-Dextran uptake which phenocopies *dOCRL^KO^*(Fig 6C). Lastly, we found increased lysotracker staining in *dOCRL^N-KO^* compared to controls and like that seen in *dOCRL^KO^* (Fig 6D). These results strongly suggest a cell-autonomous role for *dOCRL* in nephrocyte function and the underlying sub-cellular processes.

We also monitored growth and development in *dOCRL^N-KO^*animals. *dOCRL^N-KO^* larvae showed slower larval development, there was a delay of ca. 22 h for 50% of animals to complete development and pupariate (Fig 6E) and the weight and size of *dOCRL^N-KO^* 3^rd^ instar larvae was only modestly lower than that of controls (Fig 6F, Sup Fig 2 B). *dOCRL^N-KO^* larvae completed larval development, underwent pupal metamorphosis and eclosed as adults. However, only ca. 50 % of flies eclosed and the remaining did not complete pupal metamorphosis (Sup Fig 2 C, D).

### Reconstitution with *hOCRL* can rescue the phenotypes of *dOCRL^KO^*

We tested if wild type human *OCRL* (*hOCRL*) could rescue the defects observed in *dOCRL^KO^*. For this we generated transgenic flies expressing hOCRL (*HA::hOCRL*). We also generated an equivalent phosphatase dead transgene (*HA::hOCRL^PD^*) and a transgene carrying a recently reported LS-patient mutation (Ahmed P et al., 2021) with a nonsense mutation at 688 position of *hOCRL* (*HA::hOCRL*-688^C>T^). When expressed ubiquitously using hs-GAL4, both *hOCRL* and *HA::hOCRL^PD^*transgenes express a protein of the expected size, ca. 110 kDa although *HA::hOCRL^PD^*was expressed at lower levels (Fig 7B); the patient derived mutant transgene (*HA::hOCRL*-688^C>T^) expressed a truncated protein of the expected size (Fig 7B). When these transgenes were expressed in *dOCRL^KO^* using a ubiquitous promoter, *HA::hOCRL* was able to rescue the growth defect of the mutant but *HA::hOCRL^PD^* and the transgene *HA::hOCRL*-688^C>T^ (Fig 7 A) were not able to do so. Similarly, *HA::hOCRL* was able to rescue the AgNO_3_ clearance defect in *dOCRL^KO^* whereas *HA::hOCRL^PD^* and *HA::hOCRL*-688^C>T^ were unable to do so (Fig 7D). Likewise, *HA::hOCRL* was able to rescue the fluid phase uptake defect of *dOCRL^KO^* but *HA::hOCRL^PD^* and *HA::hOCRL*-688^C>T^ (Fig 7C) were unable to do so. Thus, the human ortholog rescues the defects arising from the loss of *dOCRL* and patient derived mutations can be functionally studied in the fly model.

**Figure 7:**
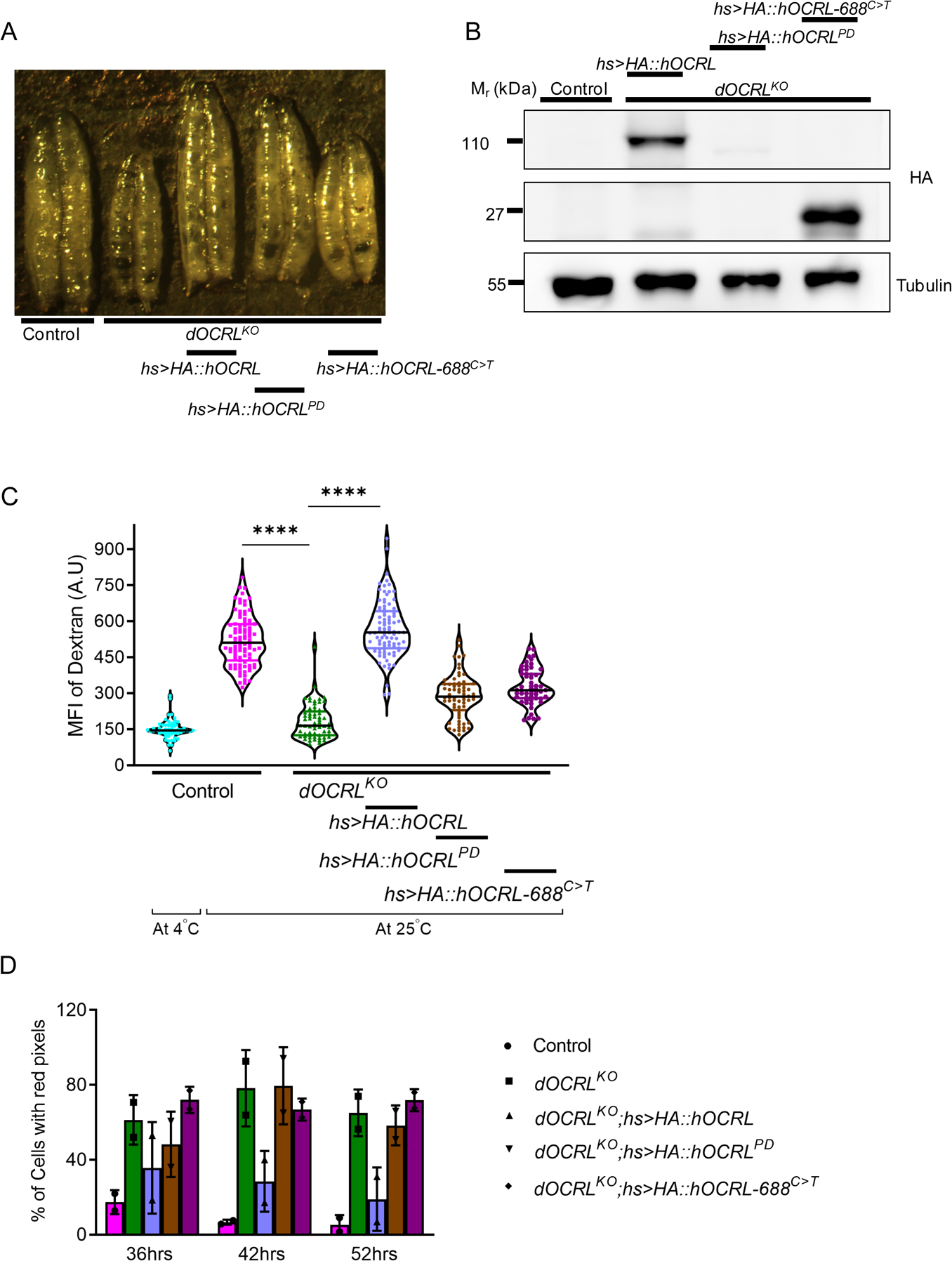
*hOCRL* reconstitution rescued the phenotypes of *dOCRL^KO^*. (A) Micrographs of third instar wandering larvae after reconstituting with the wild type *hOCRL* gene, *hOCRL^PD^*and *hOCRL*-688^C>T^. (B) Immunoblots depicting reconstitution of the wild type and transgenes with mutations in the *dOCRL^KO^*background. (C) Mean fluorescence intensity of 10KDa TMR-Dextran in nephrocytes of control, *dOCRL^KO^*, *dOCRL^KO^;hs>HA::hOCRL, dOCRL^KO^;hs>HA:: hOCRL^PD^* and *dOCRL^KO^;hs>HA::hOCRL*-688^C>T^ was calculated using ImageJ. (D) Percentage of cells with red pixel intensity corresponding to highest levels of AgNO_3_ in nephrocytes of control, *dOCRL^KO^*and *dOCRL^KO^;hs>HA::dOCRL* at 36,42 and 56 hours after transfer. Statistical tests: XY plots with mean ± S.E.M are represented. (B, D) Two tailed unpaired t-test is used. ns-Non significance, ****p<0.0001, ***p<0.001. (E) Two-way ANOVA grouped analysis with Bonferroni’s post multiple comparison tests was performed using graph pad prism to compare between each group. ns-Non significance, ****p<0.0001, ***p<0.001, *p<0.05. This statistical significance is represented on graph.

## Discussion

Clinical phenotypes in human diseases arise from altered cellular and molecular processes within cells leading to perturbed physiological processes within organ systems. Thus, when modelling human diseases, it is necessary to understand both altered sub-cellular events as well as relate these to pathophysiological changes leading to clinical phenotypes. In the case of LS, proximal tubular dysfunction is a key clinical feature and understanding its pathogenesis requires a disease model that allows the multiple sub-cellular defects ascribed to OCRL depletion to be linked to defective physiology in tubular cells. Here we describe a *Drosophila* model of LS with the following key features: (i) recapitulates the biochemical and subcellular defects previously reported for LS in human cell lines, an important feature of proximal tubule cell function in the human kidney (ii) allows the analysis of endo-lysosomal system function (iii) assess the impact of nephrocyte function in intact animals using physiological assays (iv) test *in vivo*, the pathogenic potential of patient derived LS variants by determining its impact on nephrocyte function.

During this study, using a germ line null allele of *dOCRL* (Trivedi et al., 2020) we were able to demonstrate multiple structural and functional defects in the endolysosomal system of larval nephrocytes. Several of these alterations in the early endosomal system, lysosomes, autophagosomes and the Golgi complex have been previously described in mammalian cell culture models of LS (Ungewickell et al., 2004; Vicinanza et al., 2011; Zhang et al., 1995) as well as a zebrafish larval model of LS (Oltrabella et al., 2015).This overlap of sub-cellular defects between our *Drosophila* nephrocyte model of LS and that seen in other model systems confirms that the regulation of endo-lysosomal function is a conserved feature of OCRL across multiple model systems and supports the use of *Drosophila* nephrocytes as a model for understanding the kidney deficits in LS. Moreover, it is well established that the receptors cubilin and amnionless are required for reabsorption of proteins via proximal tubular cells. The presence of cubilin and amnionless exclusively on *Drosophila* nephrocytes and their conserved function in protein uptake suggests evolutionary conservation between these cell types (Koehler and Huber, 2023). Therefore, *Drosophila* nephrocytes can provide insights into the proximal tubular dysfunction observed in LS.

It has previously been reported that in *Drosophila* nephrocytes, the handling of trace metals such as AgNO_3_ requires vesicular transport (Fu et al., 2017). Given the endolysosomal defects we noted in *dOCRL^KO^*, we tested and found defective AgNO_3_ clearance in these animals as well as enhanced sensitivity to growth on medium containing AgNO_3_. Importantly, we were able to rescue both the endocytic defects as well as the AgNO_3_ clearance delay by reconstituting *dOCRL^KO^* with a human OCRL transgene. Therefore, the sub cellular and physiological defects seen in *dOCRL^KO^*are well correlated.

We found that at the level of individual *dOCRL^KO^* nephrocytes, levels of PIP_2_, the substrate of dOCRL was elevated whereas the levels of PI4P, the product were reduced. Thus, our nephrocyte model of LS reveals the biochemical abnormality arising from loss of OCRL function. Consistent with this observation we were able to rescue both the endocytic uptake defect as well as the AgNO_3_ clearance defect by the expression of a wild type human transgene but not a phosphatase dead version of OCRL. This finding confirms that the catalytic activity of OCRL underlies the cell and whole animal physiology defects noted in our model. PIP_2_ and PI4P are both lipids implicated in the activity of many proteins involved in the regulation of endocytic trafficking thus linking the activity of OCRL to the regulation of the endosomal system, nephrocyte physiology and animal function. We also noted that nephrocyte specific deletion of dOCRL (*dOCRL^N-KO^*) was sufficient to recapitulate the altered lipid levels, endocytic defect and AgNO_3_ clearance defect seen in the germ line knockout. These finding imply a cell autonomous function for dOCRL within nephrocytes and suggest that the kidney phenotype in human patients most likely arises from the requirement of OCRL within proximal tubular cells rather than through non-cell autonomous mechanisms. This insight is particularly important and implies that future kidney specific therapeutic interventions in LS patients that may include genetic interventions targeted to proximal tubular cells are likely to be of therapeutic benefit in severe or advanced case of LS. Overall, our work provides a model system in which the cellular pathophysiology of LS can be studied with a view to designing future therapeutic strategies.

## Materials and methods

### *Drosophila* culture and strains

All flies used in this study were reared at 25 ^0^C and 50% relative humidity on fly food containing cornmeal, yeast, agar, dextrose with antibacterial and antifungal reagents in a laboratory incubator. All experiments were performed with third instar wandering larvae grown on yeast paste layered on 1% bacteriological agar. Red Oregon-R (ROR) was used as a wild type control. The following transgenic lines were obtained from Bloomington *Drosophila* stock center: Dot-GAL4 (BL6903), Dot-GAL4 (BL67609), UAS-Lifeact::RFP (BL58362), UAS-mCherry::Atg8a (BL37750), X-duplication (BL-31454); UAS-dOCRL::GFP (Avital Rodal lab), UAS-mCherry::2XFYVE (Amy Kiger, UCSD) UAS-PH-PLCδ::mCherry (Patrick Verstreken lab). UAS-P4M::GFP generated in the Padinjat lab has been previously described (Balakrishnan et al., 2018). gRNAP2dual-CG3573V, UAS-Cas9-T2A-eGFP used to generate nephrocyte specific knockout has been previously described (Trivedi et al., 2020). UAS-LTS::mCherry was generated as mentioned in (Ghosh et al., 2023).

### Generation of transgenic flies

The transgenic strains *HA::dOCRL*, *HA::hOCRL*, *HA::hOCRL^PD^, HA::hOCRL-688^C>T^* were generated by cloning cDNA into pUAST-attB with HA tag. HA::hOCRL cDNA was amplified from the pcDNA3-HA::hOCRL (Addgene-Plasmid# 22207). Not1 and Xba1 restriction sites were used to amplify the amplicon of 2730 bp and ligated into pUAST-attB vector (Drosophila Genomic Resource Center-Stock#1419). Site directed mutagenesis was used to introduce phosphatase dead mutation D523G and 688C-T mutation in pUAS-HA::hOCRL.

Oligonucleotides used:

**pUAS-HA::hOCRL:**

Not1-hOCRL-FP: GCTGCGGCCGCATGTACCCATACGACGTC

Xba1-hOCRL-RP: GCTTCTAGATTAGTCTTCTTCGCT

**pUAS-HA::hOCRL688^C-T^:**

hOCRL688^C-T^-FP: ATCCTGGCAAAGTGAGAGAAAGAATA

hOCRL688^C-T^-RP: ATTCTTTCTCTCACTTTGCCAGGATA

**pUAS-HA::hOCRL-PD:**

hOCRL^D523G^-FP: CTGAAAACCAGCGGCCACAAGCCTGT

hOCRL^D523G^-RP: ACAGGCTTGTGGCCGCTGGTTTTCAG

### Quantitative RT-PCR

Total RNA was extracted from third instar wandering larvae using TRIzol reagent (Invitrogen) followed by treatment with DNase-I (Invitrogen). cDNA was synthesized using Superscript II RNase H-Reverse transcriptase (Invitrogen) and random hexamers (Applied Biosystems). Non-template and no reverse transcription control samples were also included. Quantitative PCR was performed using Applied Biosystem 7500 Fast Real-Time PCR system using cDNA samples so synthesized. In addition to the transcripts of interest, Ribosomal Protein 49(*RP49*) was amplified as a housekeeping gene. Both *dOCRL* and *RP4*9 primers were designed at exon-exon junction with primers attaining all the parameters recommended for q-PCR primer designing. The C_t_ values obtained for each gene was normalized with the values of RP49 of the sample. Biological triplicates were assayed for each genotype.

The primers used for q-PCR were:

RP49 forward: CGGATCGATATGCTAAGCTGT

RP49 reverse: GCGCTTGTTCGATCCGTA

dOCRL forward: GAACAACAAGACCTGCAGC

dOCRL Reverse: CTGTCCATCATCTTATCGATCC

### Growth curve assay

In this assay *wildtype* and *dOCRL^KO^* flies were allowed to lay eggs for 4-6 h on normal fly food. 24 h post egg laying, first instar larvae were carefully transferred onto yeast media with controlled crowding. Around 20 larvae was transferred per vial containing yeast media layered on agar, 6 biological replicates were maintained and allowed to grow at 25 ^0^C. After 96 hrs, the number of larvae pupariated was noted for both wildtype and *dOCRL^KO^* in 3 h time intervals. The number of larvae pupariated were plotted to give the percentage of pupariation achieved. The mean pupariation percentage were calculated from each time bin. These pupariation percentage was plotted and fitted to variable slope on Graph Pad Prism as shown in Fig 1E.

### Nephrocyte function assay

Flies were allowed to lay eggs on normal food. 24 h after egg laying, first instar larvae were transferred to yeast paste containing 0.003% AgNO_3_ layered on 1% bacteriological agar. After 32 h of feeding on AgNO_3_ media, larvae were transferred to yeast paste media without AgNO_3_. 36, 42 and 56 h post transfer from AgNO_3_ media, larvae were dissected, fixed with 4% paraformaldehyde for 15 minutes at room temperature, washed thrice using PBS, mounted in 70% glycerol. Preparations were subjected to bright field imaging under a 10X objective using a Olympus BX43 microscope and digital images were recorded using CellSens software. These images were converted to 8-bit 16 color pixels corresponding to the intensity of AgNO_3_, from white being highest concentration to black being least concentration. The number of cells with red pixels and yellow pixel, corresponding to high and low concentration of AgNO_3_, was calculated and plotted. The experiment was repeated in three individual trials with triplicates; 60 nephrocytes were used for quantification in each genotype. For survival analysis, first instar larvae were transferred to media containing various concentrations of AgNO_3_ media and the percentage larval survival was calculated.

### Dextran uptake assay

Wandering third instar larvae were washed thrice with PBS. Nephrocytes were dissected in Brodie and Bate’s buffer (B&B buffer) (consists of 135 mM NaCl, 5 mM KCl, 4mM MgCl_2_, 2mM CaCl_2_, 5mM TES, 36mM sucrose). For dextran uptake by nephrocytes, 0.33mg/ml of TMR-dextran (Thermo-fisher scientific, # D1868) was pulsed for 5 minutes in B&B buffer at 25^0^C in dark. As negative control another set was incubated in dextran at 4^0^C. At the end of the incubation period, preparations were washed for 10 minutes in ice-cold PBS, fixed in 4% paraformaldehyde (PFA) in PBS for 15 minutes at room temperature (25^0^C) and washed thrice with PBS. The tissues were mounted on glass slide with 70% glycerol in PBS. Nephrocytes were imaged on an Olympus FV3000 laser scanning confocal microscope and Fiji ImageJ was used for quantification of mean fluorescence dextran intensity per unit area of nephrocytes from the maximum projection images. These images were used to estimate the area of nephrocytes by marking the cell boundary using ImageJ. The assay was repeated thrice with three larvae in each trial.

### mBSA uptake assay

mBSA was synthesized and conjugated with Cy5 bis-functional dye (Cy5 Bis NHS Ester, Cat no. C183, GeneCopoeia) according to manufacturer’s instructions. Nephrocytes were dissected as described above and immediately incubated with conjugated mBSA (0.1mg/ml) for 15 minutes. Subsequently, cells were washed, fixed with 4% PFA for 15 minutes. Cells were washed, stained with DAPI and imaged on an Olympus FV3000 confocal microscope. Mean fluorescence intensity (MFI) of mBSA was quantified using ImageJ. The assay was repeated three times with three larvae in each trial.

### Western Blot

All blots were performed from third instar wandering larval lysates except for those where probe levels were estimated; in this case, larval nephrocytes were dissected in PBS and processed for lysate extraction. Lysates were prepared by homogenizing third instar wandering larvae or dissected larvae with Tris-Lysis buffer containing Roche Protease inhibitor and PhosStop and heated at 95^°^C for 5 minutes. Samples were separated by SDS-Polyacrylamide gel electrophoresis and transferred onto nitrocellulose filter membrane [Hybond-C Extra; (GE Healthcare, Buckinghamshire, UK)] using wet transfer (BioRad, California, USA). Membranes were blocked with 5% Blotto (sc-2325, Santa Cruz Biotechnology, Texas, USA) in PBS with 0.1% Tween 20 (Sigma Aldrich) (PBST) for 2 h at room temperature (25 ^°^C) followed by incubation in primary antibody diluted in 5% Blotto in PBST overnight at 4 ^0^C. Primary antibodies dilutions used were 1:1000 anti-dOCRL (Avital Rodal lab), 1:1000 anti-HA (CST, #2367S), 1:4000 anti-α-tubulin-E7c (DHSB, lowa, USA), 1:1000 anti-Syntaxin1A (DSHB, Iowa, USA), 1:3000 anti-RFP (Rockland, 600-401-600-RTU), 1:1000 anti-α-actin (Sigma Aldrich), 1:1000 anti-mcherry (Thermo, PA5-34974) 1:2000 anti-GFP (Santa cruz, sc9996). Following this, the membrane was washed for 10 minutes thrice with PBST and incubated with the appropriate HRP conjugated secondary antibody (Jackson Laboratories, USA) at 1:10000 dilution in PBST for 2 hours at room temperature. After this, the membrane was washed for 10 minutes thrice with PBST. Blots were developed with ECL substrate (GE healthcare) and imaged using GE Image quant LAS 4000 system. The blots were stripped by incubating with 3% glacial acetic acid in PBST for 30 minutes, washed thrice and re-probed with primary antibody. Quantification of signals was done using Fiji ImageJ software and the values were normalized to the loading control intensity.

### Immunostaining of nephrocytes

Wandering third instar larvae were dissected in Brodie & Bate’s buffer and fixed with 4% PFA in PBS for 15 minutes. Cells were permeabilized with 0.3% PBSTX (0.3% Triton X in PBS) for 30 minutes three times and blocked with 5% FBS in PBSTX for 2 h at room temperature. Primary antibodies were diluted with blocking solutions at incubation was done at 4^0^C overnight. The following primary antibodies were used: anti-dOCRL 1:200 (Avitol Rodal lab), anti-Rab7 1: 200 (DSHB), anti-EEA1 1:800 (Marino Zerial, MPI-CBG, Dresden, Germany). Larval tissues were then washed with PBSTX at room temperature three times with 30 minutes, incubated with appropriate secondary antibody conjugated with fluorophore at 1:300 dilution in blocking solution for 3 h at room temperature. Tissues were washed with PBSTX and mounted on glass slides with 70% glycerol in PBS. Samples were imaged on an Olympus FV3000 laser scanning confocal microscope and Fiji ImageJ was used for quantification of mean fluorescence intensity per unit area of nephrocytes. MFI of EEA-1 and Rab-7 staining was quantified after background subtraction from the maximum projections of the stacks and normalized to the area of nephrocytes.

### Lysotracker Assay

Wandering third instar larvae was dissected in B&B buffer rapidly and incubated with Lysotracker deep red (Thermo-fisher scientific) at 1:1000 dilution in B&B buffer for 5 minutes at room temperature. Following this, tissues were washed thrice in ice cold PBS and fixed with 4% PFA in PBS for 4 minutes (mild fixation as harsh fixation might lead to bleaching of the dye) and mounted on glass slide with 70% glycerol in PBS. Samples were imaged immediately on an Olympus FV3000 laser scanning confocal microscope and Fiji ImageJ was used for quantification of mean fluorescence intensity per unit area of nephrocytes.

### DNA extraction for verification of *dOCRL^N-KO^*

Around 8 control and *dOCRL^N-KO^* third instar wandering larvae was dissected for nephrocytes in B&B buffer. The buffer was drained from the dissected tissue and carcass transferred into a 0.5 ml micro centrifuge tube. Samples were homogenized in 50µl of squishing buffer containing 200µg/ml proteinase K using a pipette tip and incubated for 30 minutes at room temperature (25 ^0^C). Following this Proteinase K was inactivated by heating at 95 ^0^C for 5 minutes. 3µl of this lysate was used for verification of *dOCRL^N-KO^* by PCR analysis. We used CG7004 gene as a control to determine the quality of DNA sample preparation.

### PIP and PIP_2_ measurement from larvae by LC-MS/MS

#### Lipid isolation for PIP and PIP_2_ measurements

Lipids were isolated from three third instar wandering larvae. Total lipids were isolated and neomycin chromatography was performed as described earlier(Ghosh et al., 2019). 25 ng each of 17:0 | 20: 4 PI4P (Cat# Avanti LM 1901) and 17:0 | 20: 4 PI(4,5)P_2_ (Cat# Avanti LM 1904) were used as internal standards and added prior to homogenization of larvae during extraction.

#### Total Organic Phosphate measurement

600 µl flow-through obtained from the phosphoinositide binding step of neomycin chromatography was obtained from the last step of lipid extraction and stored separately in phosphate free glass tubes till assay was performed. The sample was heated in a dry heat bath at 90^0^C in phosphate-free glass tubes until it dried (Cat# 14-962-26F). The rest of the process was followed as described previously (Jones et al., 2013).

#### LC-MS/MS

The instrument operation was as described in our previous work on PI5P quantification (Ghosh et al., 2019) (Ghosh and Raghu, 2021). For *in vivo* lipid measurements, the samples were washed with post-derivatization step before injecting in mass spec. Samples were run on a hybrid triple quadrupole mass spectrometer (Sciex 6500 Q-Trap) connected to a Waters Acquity UPLC I class system. Separation was performed on a ACQUITY UPLC Protein BEH C4, 300Å, 1.7 µm, 1 mm X 100 mm column [Product #186005590] using a 45% to 100% acetonitrile in water (with 0.1% formic acid) gradient over 10 mins. MS/MS and LC conditions used were as described earlier (Ghosh et al., 2019).

#### Software and data analysis

Image analysis was performed by Fiji software (Open source). Mass spec data was acquired on Analyst® 1.6.2 software followed by data processing and visualization using MultiQuant^TM^ 3.0.1 software.

#### MS parameters and List of MRMs

MS was run in positive mode. Dwell time of 65 milliseconds were used for experiments with CAD value at medium range, GS1 and GS2 at 20, CUR (Curtain gas) at 37, IS (ESI Voltage) as 5200 and TEM (Source Temperature) as 300, DP (Declustering Potential) at 140 for PIPs, 60 for PIP_2_s, EP (Entrance Potential) at 12 for PIPs, 11 for PIP_2_s, CE (Collision Energy) at 29 for PIPs, 37 for PIP_2_s, CXP (Collision cell Exit Potential) at 15 for PIPs, 12 for PIP_2_s. Both Q1 and Q3 masses were scanned at unit mass resolution.

| Acyl chain length | PIP parent ion | PIP<sub>2</sub> parent ion | Daughter ion 496 |
| --- | --- | --- | --- |
| 30:1 | - | 1011.5 | 521.5 |
| 32:0 | 933.5 | 1041.5 | 551.5 |
| 32:1 | 931.5 | 1039.5 | 549.5 |
| 32:2 | 929.5 | 1037.5 | 547.5 |
| 34:0 | 961.5 | 1069.5 | 579.5 |
| 34:1 | 959.5 | 1067.5 | 577.5 |
| 34:2 | 957.5 | 1065.5 | 575.5 |
| 34:3 | 955.5 | 1063.5 | 573.5 |
| 34:4 | - | 1061.5 | 571.5 |
| 36:0 | - | 1097.5 | 607.5 |
| 36:1 | 987.5 | 1095.5 | 605.5 |
| 36:2 | 985.5 | 1093.5 | 603.5 |
| 36:3 | 983.5 | 1091.5 | 601.5 |
| 36:4 | 981.5 | 1089.5 | 599.5 |
| 38:0 | - | 1125.5 | 635.5 |
| 38:1 | - | 1123.5 | 633.5 |
| 38:2 | - | 1121.5 | 631.5 |
| 17:0\| 20:4 ISD | 995.5 | 1103.5 | 613.5 |

### Statistical Analysis

Raw data of imaging were processed and analyzed using Fiji ImageJ, processed data was plotted, and the statistical analysis were analyzed using GraphPad Prism. For all experiments, error bars represent SEM, and p values were calculated by using the student’s unpaired t-test with welch correction. * P < 0.05, ** P < 0.01, *** P < 0.001. To establish significance in growth curve analyses and nephrocyte filtration assay a two-way Annova was performed. Survival analysis was done for survival assay.

## Acknowledgements

This work was supported by the Department of Atomic Energy, Government of India, under Project Identification No. RTI 4006, a Wellcome-DBT India Alliance Senior Fellowship to PR (IA/S/14/2/501540) and a Department of Biotechnology, Government of India grant to AV (BT/PR11030/MED/30/1644/2016).

## Figure Legends

**Supplementary Figure 1:**

Immunoblots to verify the expression of Ph-PLCδ::mCherry (A), P4M::GFP (C), mCherry::2XFYVE (E), LTS::mCherry (G) and mCherry::ATG8a (I) probe levels in control and *dOCRL^KO^*nephrocytes. Quantification of probe levels of Ph-PLCδ::mCherry (B), P4M::GFP (D), mCherry::2XFYVE (F), LTS::mCherry (H) and mCherry::Atg8a (J). Statistical tests: XY plots with mean ± S.E.M are represented. Two tailed unpaired t-test is used. ***p<0.001, *-p<0.05.

**Supplementary Figure 2: Nephrocyte specific *dOCRL* knockout phenocopies the defects of whole body *dOCRL^KO^***

(A) Verification of knock out of dOCRL gene in nephrocytes by agarose gel electrophoresis in control, *dOCRL^KO^* and *dOCRL^N-KO^*. (B) Micrographs of third instar larvae of control, *dOCRL^N-KO^* and *dOCRL^KO^*. (C-D) Percentage of control and *dOCRL^N-KO^* larvae pupariated and the percentage of adult flies eclosed respectively. (E) Confocal micrographs of PIP_2_ localization in control, *dOCRL^KO^* and *dOCRL^N-KO^*. (F) Quantification of plasma membrane to cytoplasmic MFI of PIP_2_ measured using ImageJ. Statistical tests: XY plots with mean ± S.E.M are represented. Two tailed unpaired t-test is used. ***p<0.001, *-p<0.05.

