## Supplementary figures and images for "A genetic and physiological model of renal dysfunction in Lowe syndrome"

### supplementary figures

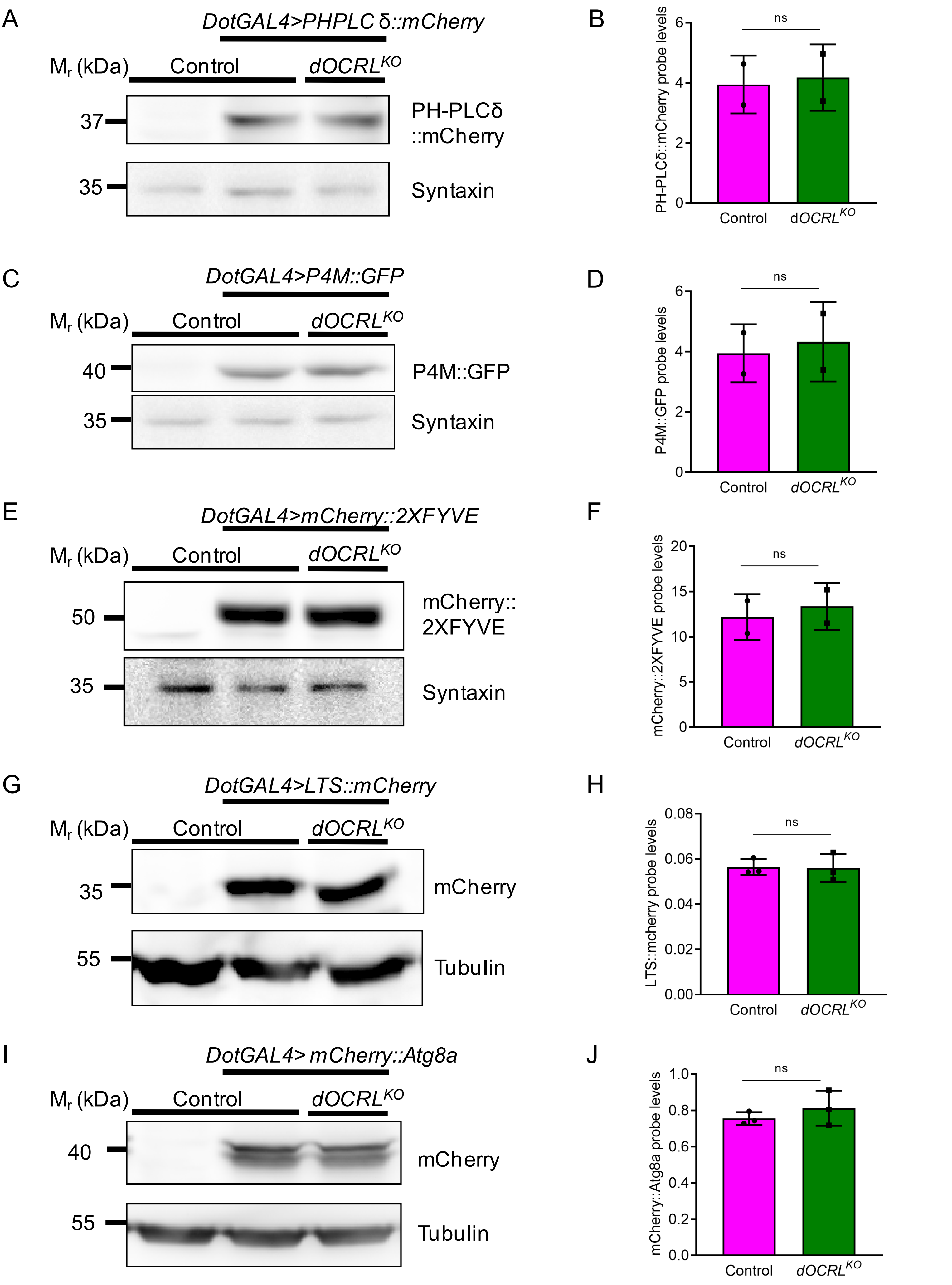


**Ramesh et.al, supplementary figure 1**

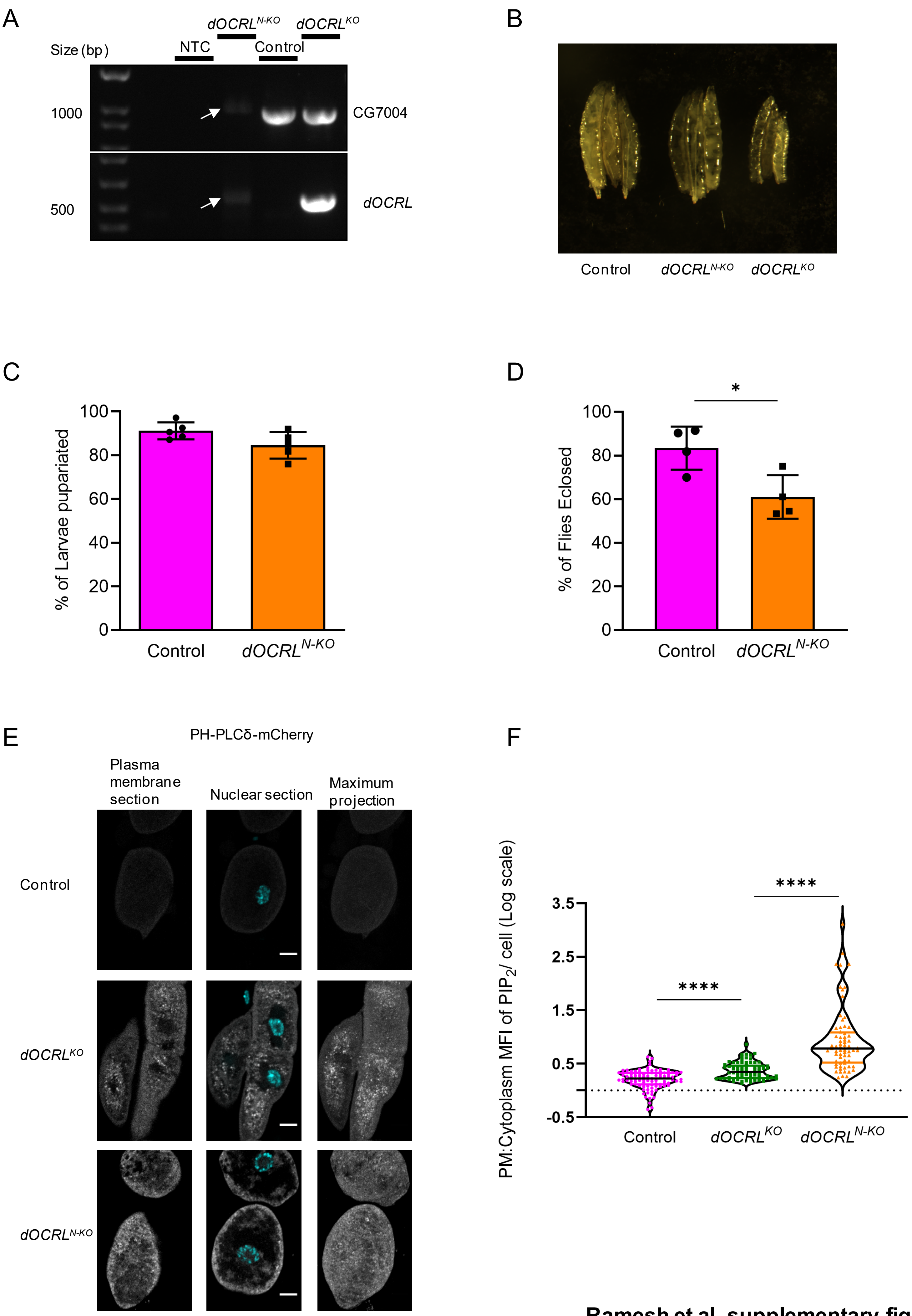


**Ramesh et.al, supplementary figure 2**
